# Utility of recombinant envelope domain III as a diagnostic antigen for the specific detection of Kyasanur Forest Disease

**DOI:** 10.1101/2025.09.07.674709

**Authors:** Sayad Hafeez, Rajeshwara Achur, Easwaran Sreekumar, Asha Srinivasan, Suchetha Kumari, N. B. Thippeswamy

## Abstract

Kyasanur Forest Disease (KFD), commonly known as “monkey fever,” is a highly neglected tropical disease caused by the Kyasanur Forest Disease Virus (KFDV). KFD is endemic to Western Ghats of Karnataka, India, with seasonal outbreaks during December to June every year. As there is no standard treatment regime, KFD can be fatal with a mortality rate of 2-10%. Currently, KFD is detected through a non-specific IgM-ELISA followed by RT-PCR, which often delays diagnosis, leading to increased disease severity and even death. To address this, we focused on developing a specific antigen-based KFD detection. The KFDV Envelope Domain III (EDIII) and Non-Structural 1 (NS1) proteins were chosen as detection markers, cloned, and expressed using pET28a(+) vector in BL-21 (Rosetta) *E. coli* and purified. These proteins were used to raise polyclonal antibodies in rabbits and the antibody titre was found to be 1:256,000 and 1:512,000 against rEDIII and rNS1 proteins, respectively. Importantly, these polyclonal antibodies showed no cross-reactivity against corresponding dengue virus EDIII and NS1 proteins. Using polyclonal antibodies against rEDIII, we developed sandwich ELISA for the specific detection of KFD, which has demonstrated high specificity and sensitivity. Further, anti-rEDIII polyclonal antibodies also detected full-length KFDV-E protein expressed in mammalian cells, confirming the antibody specificity for the native viral antigen.

## 1. Introduction

Kyasanur Forest Disease (KFD), a highly neglected viral tropical disease of zoonotic origin, with a seasonal outbreak during December to June, is caused by the Kyasanur Forest Disease Virus (KFDV) (1). The virus is transmitted by the bite of ticks, *Haemophysalis spinigera*, which serves both as a vector and disease reservoir (2). KFDV belongs to the family Flaviviridae having 11 kb positive-sense RNA genome and 45 nm diameter. The genome codes for three structural proteins (capsid, precursor membrane, and envelope) and seven non-structural proteins (NS1, NS2A, NS2B, NS3, NS4A, NS4B, and NS5) (3). KFD has accounted for about 823 confirmed cases and 50 deaths during 2003-2012 out of ∼5000 suspected cases. KFD still accounts for about 100-500 cases every year in the endemic zone holding a mortality rate of ∼2-10% (4,5). During 2018-2024, the disease severity has taken a death toll of ∼35-40 and shown its presence in the adjacent states such as Kerala, Maharashtra, Goa and Gujarat (6,7).

Currently, there is no effective treatment for KFD and the formalin-inactivated vaccine used, has become less effective (8). Further, the rapid KFD diagnosis remains a challenge and currently being detected by IgM-ELISA and RT-PCR (nested and real-time) by targeting KFDV-NS5 gene (9). The IgM-ELISA focuses on antibodies generated after few days of infection, which delays disease detection, leading to severe illness of infected individuals. Further, this method requires validation by RT-PCR due to potential cross-reactivity with dengue virus (DENV). Since KFDV is classified as a BSL-4 pathogen, RT-PCR must be conducted in specialized laboratories by trained personnel, and is time-consuming. Despite being identified nearly six decades ago, there is no rapid and specific diagnostic test for KFD. Thus, there is an urgent need to develop early KFD diagnostics to alleviate both health risks and the disease associated economic burden.

Studies on flaviviruses indicate that the Envelope (E) and non-structural (NS) proteins are valuable targets for diagnosis and vaccine development. The flaviviral E protein is antigenic and is involved in viral adherence, penetration, hemagglutination, and acts as a fusion peptide. The E protein has three domains, among these, envelope domain III is a diagnosis marker target containing distinct virus-specific epitopes which avoids cross-reactivity (10). NS3 exhibits several enzymatic functions, including RNA helicase and protease activities, which is involved in polyprotein cleavage. The 146 to 154 amino acid sequence of NS3 is highly conserved among flaviviruses which serves as a positive marker for cross-reactivity (11). The NS5 is the largest non-structural protein which plays a crucial role in replication having RNA polymerase and methyltransferase activities, in addition to involving in *de novo* RNA synthesis (12).

NS1 exhibits multiple functions, its interaction with NS4A-4B stabilizes the transmembrane proteins in the lumen of endoplasmic reticulum facilitating viral replication (13). The NS1 exists in different forms, including as a secretory variant released by infected host cells into the bloodstream. The secreted NS1 also plays a key role in undermining host immune mechanisms, particularly the innate immune system (14). The NS1 based ELISA detection of DENV has been a remarkable success in the development of early detection kit. Currently, various strip-based immunochromatographic kits are also available for dengue detection (15,16). The efficacy of dengue diagnosis has improved with the introduction of NS1 antigen and anti-DENV specific IgM/IgG antibodies into strips, aiding in the interpretation of infection stages (17). Given these characteristics and the immunodominance of envelope and NS1 proteins, we employed these to develop an antigen-based KFD rapid diagnostic kit.

## 2. Materials and methods

### 2.1 Experimental ethics

8 to 10-week old female New Zealand rabbits weighing ∼2.5 to 3 kg were used for raising polyclonal antibody. 6 to 8-weeks old BALB/c mice weighing ∼22-25 g were used for capture antibody production. All the animals were maintained in 12h light/dark cycle with water ad libitum as per the OECD guidelines and protocols approved by the IAEC ***(Protocol number: CBPL-IAEC-031/06/2023)***. Uninfected human serum samples and dengue positive samples collected and stored at K.S. Hegde Medical Academy, Mangaluru, were used for our analysis ***(Ethics statement: EC/NEW/INST/2022/KA/0174)***.

### 2.2 Bacterial Expression and Purification of Recombinant KFDV-EDIII and NS1 Proteins

The amino acid sequences of KFDV EDIII and NS1 proteins were retrieved from UniProt and back-translated to nucleotide sequence (nBLAST, NCBI). Codon-optimized sequences for *E. coli* were synthesized (EDIII by GeneScript, USA; NS1 by Twist Biosciences, USA) and cloned into the pET28a(+) vector at NdeI/XhoI sites. Plasmids were first transformed into *E. coli* DH5-α for amplification, extracted using alkaline lysis and then transformed into *E. coli* BL21-Rosetta for protein expression ***(IBSC number: JSSAHER/IBSC-006/2021)***. A single colony was cultured in LB medium with kanamycin (50□μg/ml) and chloramphenicol (34□μg/ml), first in 10 ml (overnight), then scaled to 500 ml. Cultures were induced with 0.5□mM IPTG and incubated for 4h at 37°C until OD□□□ of ∼0.6–0.8 is attained. Cells were harvested by centrifugation (5,000 rpm, 15□min), resuspended in native buffer (20□mM Tris, 300□mM NaCl, 20% glycerol, pH□8.0), and sonicated on ice (50% amplitude, 5□min on/3□min off). Lysates were centrifuged (5,000 rpm, 20□min), and pellets solubilized in denaturing buffer containing 8□M urea, at 37°C for 1h with shaking. After centrifugation (12,000 rpm, 30□min), the supernatant was loaded onto Ni-NTA resin equilibrated with denaturing buffer, washed with buffer containing 30□mM imidazole, and the proteins were eluted with 300□mM imidazole. Eluted fractions were pooled and refolded by stepwise dialysis against native buffer with decreasing urea (6□M to 0□M). Protein concentration and purity were assessed using Bradford’s reagent (Biorad) and SDS-PAGE (15% for rEDIII, 12% for rNS1).

### 2.3 Polyclonal Antibody Production in Rabbits and Mice

Female New Zealand rabbits (8–10 weeks old, 2.5–3□kg, *n*=2) and BALB/c mice (6–8 weeks old, 22–25□g, *n*=6) were used for polyclonal antibody production against rEDIII and rNS1 proteins and the pre-immune serum was collected one day prior to immunization. For primary immunization, rabbits were subcutaneously injected with 500□µg/ml of rEDIII or 250□µg/ml of rNS1 in 1□ml PBS (100□mM, pH 7.4) emulsified with an equal volume of Freund’s complete adjuvant (FCA) (Invivogen). Mice were injected subcutaneously with 50□µg/ml rEDIII or 30□µg/ml rNS1 in 100□µl PBS with an equal volume of FCA. For all the animals, two booster doses were administered at 21-day intervals using half the initial antigen dose with Freund’s incomplete adjuvant. The first test bleed was collected after 14 days of initial immunization, with subsequent bleeds at 8 days post each booster dose (Fig. 2A).

### 2.4 Indirect ELISA

The immunogenicity of rEDIII and rNS1 proteins was assessed using indirect ELISA (18). Briefly, 200□ng of each protein in 100 µl carbonate buffer (0.1□M, pH□9.4) was coated onto 96-well NUNC Maxisorp plates (Thermo Scientific) and incubated overnight at 4°C. Plates were washed with PBS (10□mM, pH□7.4) and blocked with 1% BSA in PBS (50□mM, pH□7.4) for 1h at 37°C. After washing with PBST (PBS with 0.05% Tween-20), serial dilutions (1:1,000 to 1:10,24,000) of rabbit serum were added in blocking buffer and incubated for 1h at 37°C. After 3×5 min washes with PBST, HRP-conjugated secondary antibody (Sigma, 1:40,000 in blocking buffer) was added and incubated for 1h at 37°C. TMB substrate (Sigma) was added to develop color, and the reaction was stopped with 2□M sulfuric acid and the absorbance was measured at 450□nm.

### 2.5 Sandwich ELISA

A novel sandwich ELISA was developed using mouse anti-rEDIII or rNS1 polyclonal antibodies as capture antibodies. Briefly, capture antibodies (1:50 in 0.1□M carbonate buffer, pH□9.4) were coated onto 96-well NUNC Maxisorp plates (Thermo Scientific) and incubated overnight at 4°C. Plates were washed with PBS (10□mM, pH□7.4) and blocked with 1% BSA in 50□mM PBS (pH□7.4) for 1h at 37°C. After washing with PBST (PBS with 0.05% Tween-20), serial dilutions (0.7□ng to 2.5□µg) of rEDIII and rNS1 proteins were added and incubated for 1h at 37°C. Following 3x5 min washes, 1:1000 diluted rabbit anti-rEDIII or anti-rNS1 polyclonal antibodies (in blocking buffer) were added and incubated for 1h at 37°C. The plates were washed again and incubated with HRP-conjugated secondary antibody (Sigma, 1:40,000 in blocking buffer) for 1h at 37°C. TMB substrate (Sigma) was added to develop color, and the reaction was stopped with 2□M sulfuric acid and the absorbance was measured at 450□nm.

### 2.6 Dot-Immunoblot Assay

The dot-blot assay was performed to evaluate antibody titers against rEDIII and rNS1 as per method described earlier with minor modifications (19). PVDF membranes were pre-activated by soaking in methanol (30□s), rinsed in distilled water (2□min), and equilibrated with TBST (20□mM Tris, 0.1% Tween-20, pH□7.5). The antigens, rEDIII and rNS1 were prepared (1□mg/ml in 10□mM TBS, pH□7.5), and 2–5□µg was dotted onto the membrane and air-dried. Membranes were blocked with 3% skimmed milk in TBS for 1h at RT with gentle shaking, followed by a 5□min TBST wash. The membranes were then incubated with serially diluted rabbit serum prepared in blocking buffer for 1h at RT, with intermittent shaking. After 3×10□min TBST washes, HRP-conjugated secondary antibody (Sigma, 1:20,000 in blocking buffer) was added and incubated for 1h with shaking. Color development was carried out using 1% 3-amino-9-ethylcarbazole (AEC) in *N,N*-dimethylformamide (DMF), prepared in 0.05□M acetate buffer (pH□5.5) containing 0.3% hydrogen peroxide.

### 2.7 SDS-PAGE

The expression and purity of rEDIII and rNS1 were assessed by Tris-glycine SDS-PAGE (20. Laemmli, 1970). For rEDIII, *E. coli* lysates (pre- and post-IPTG induction) and purified protein were resolved using 4% stacking and 15% separating gels at 150□V for 150□min. For rNS1, the samples as above were run on 4% stacking and 12% separating gels at 150□V for 90□min. Gels were stained with Coomassie Brilliant Blue to visualize protein bands and the protein sizes were determined by comparing with a pre-stained molecular weight marker (Himedia, 11–245□kDa).

### 2.8 Western Blotting

Proteins separated by SDS-PAGE were transferred by semidry method onto methanol-activated PVDF membranes at 25□V for 45□min (Trans-Blot, Bio-Rad). Membranes were blocked with 1% BSA in TBST (10□mM TBS, 0.05% Tween-20) for 1h at RT, followed by 3×10 min TBST washes. For expression detection, membranes were incubated for 1h with anti-His tag antibody (CST, USA; 1:1000). For cross-reactivity analysis, rabbit anti-KFDV-rEDIII and anti-rNS1 antibodies (1:1000) were used. After washing, membranes were incubated for 1h with HRP-conjugated secondary antibody (Sigma-Aldrich; 1:20,000 in blocking buffer). After 3×10 min washes, chromogenic HRP substrates, either TMB or AEC (Sigma-Aldrich) were applied for color development and the blots were imaged (GelDoc EZ imager, Bio-Rad, USA).

### 2.9 DENV sample analysis

The strip based DENV lateral flow assay kits (Agappe, India) were used to detect the DENV in infected patient samples. DENV-NS1 antigen microelisa kit (J Mitra, India) was used to verify the cross-reactivity of anti-rEDIII and rNS1 polyclonal antibodies against DENV-NS1 positive samples. Both assays were performed as per the manufacturer’s protocol. DENV rEDIII and rNS1 proteins were purchased commercially (Native antigen company, UK) for cross-reactivity analysis.

### 2.10 Expression of KFDV E Protein in Mammalian Cells and Immunoreactivity Analysis

The complete KFDV envelope coding sequence (1487□bp, based on GenBank MHD13227) was synthesized and cloned into the pcDNA3.1-hygro vector. Briefly, to prepare for transfection, HEK293 cells (3 × 10□) were seeded into 6-well plates and incubated for 18 h. The cells were then co-transfected with 2 µg/ml of pcDNA-KFDV E expression plasmids using polyethyleneimine (Sigma) at a concentration of 8 µg/ml. Stable expression of the full-length KFDV E protein was achieved by selecting the transfected cells with hygromycin (50 µg/ml) over a period of 60 days. The resulting HEK293 producer cells underwent clonal selection and were subsequently cryopreserved. For downstream applications, these producer HEK293 cells were cultured for an additional 24 h. The cells were washed with PBS and lysed using RIPA buffer supplemented with protease inhibitor cocktail (Pierce). Lysates were incubated on ice for 45□min with vertexing every 10□min, followed by centrifugation at 10,000×g for 15□min. The supernatant containing the E protein was collected and quantified using Bradford reagent. Thirty micrograms of total protein were separated by SDS-PAGE and transferred onto a PVDF membrane. Membranes were blocked with 5% BSA in TBST for 1h at RT and incubated overnight at 4°C with primary antibodies: rabbit anti-rEDIII (1:500), anti-rNS1 (1:1000), and mouse anti-β-actin (1:3000) in 3% BSA-TBST. Following 3×10 min TBST washes, membranes were incubated with HRP-conjugated secondary antibodies—anti-rabbit IgG-HRP and anti-mouse IgG-HRP (CST, USA) for 90□min. The chemiluminescent was developed by using West Pico Plus substrate (Thermo Scientific, USA), and analyzed using a pre-stained protein ladder (Thermo Scientific).

### 2.11 Statistical Analysis

All the data were analyzed using GraphPad Prism v8.3. Antibody titers were evaluated using nested one-way ANOVA followed by Sidak’s multiple comparison test. One-way ANOVA was applied to assess cross-reactivity of anti-rEDIII and rNS1 antibodies against DENV patient samples. One-way ANOVA with Dunnet’s multiple comparison was used for the sensitivity and specificity analysis of sandwich ELISA results. Significance levels were denoted as follows: *P*□<□0.05 (\**), P*□*<*□*0*.*01 (*\*\**), P*□*<*□*0*.*001 (*\*\*\*), *P*□<□0.0001 (****), and *ns* (not significant). Data were presented as mean□±□SEM.

## 3. Results

### 3.1 Bacterial expression and purification of KFDV-rEDIII and rNS1 proteins

The KFDV-EDIII and NS1 protein sequences were sourced from the European protein database, UniProt. We optimized these sequences for bacterial expression and seamlessly incorporated them into the pET28a(+) bacterial expression vector at multiple cloning sites of NdeI and XhoI (Figs. 1A and 1D). Additionally, we integrated a six-histidine tag at the N-terminal of the recombinant KFDV proteins to aid in purification. To boost the copy number, the plasmid vector was transformed into *E. coli* DH5-α strain. Subsequently, the purified plasmid was introduced into *E. coli* BL21 strain to facilitate effective protein expression. Induction was successfully achieved for rEDIII and rNS1 proteins with 0.5 mM IPTG. Ni-NTA columns were used for the purification which yielded 0.6 mg/ml (rEDIII) and 0.4 mg/ml (rNS1) proteins. The SDS-PAGE analysis indicated the size of rEDIII and rNS1 proteins to be ∼10 kDa and ∼40 kDa, respectively (Figs. 1B and 1E). Additionally, densitometry analysis showed a single peak for both purified rEDIII and rNS1 proteins, with intensities (arbitrary units) of approximately 200 and 150, respectively. This indicates that the purity of both recombinant proteins is ≥90% (Supplementary Fig. S1). To confirm the expression of rEDIII and rNS1, western blot analysis was performed, using monoclonal anti-his tag antibody (Figs. 1C and 1F). This compelling evidence indicates the successful expression of both rEDIII and rNS1 proteins in bacterial system, making these as prime candidates for immunization and subsequent production of polyclonal antibodies.

**Figure 1:**
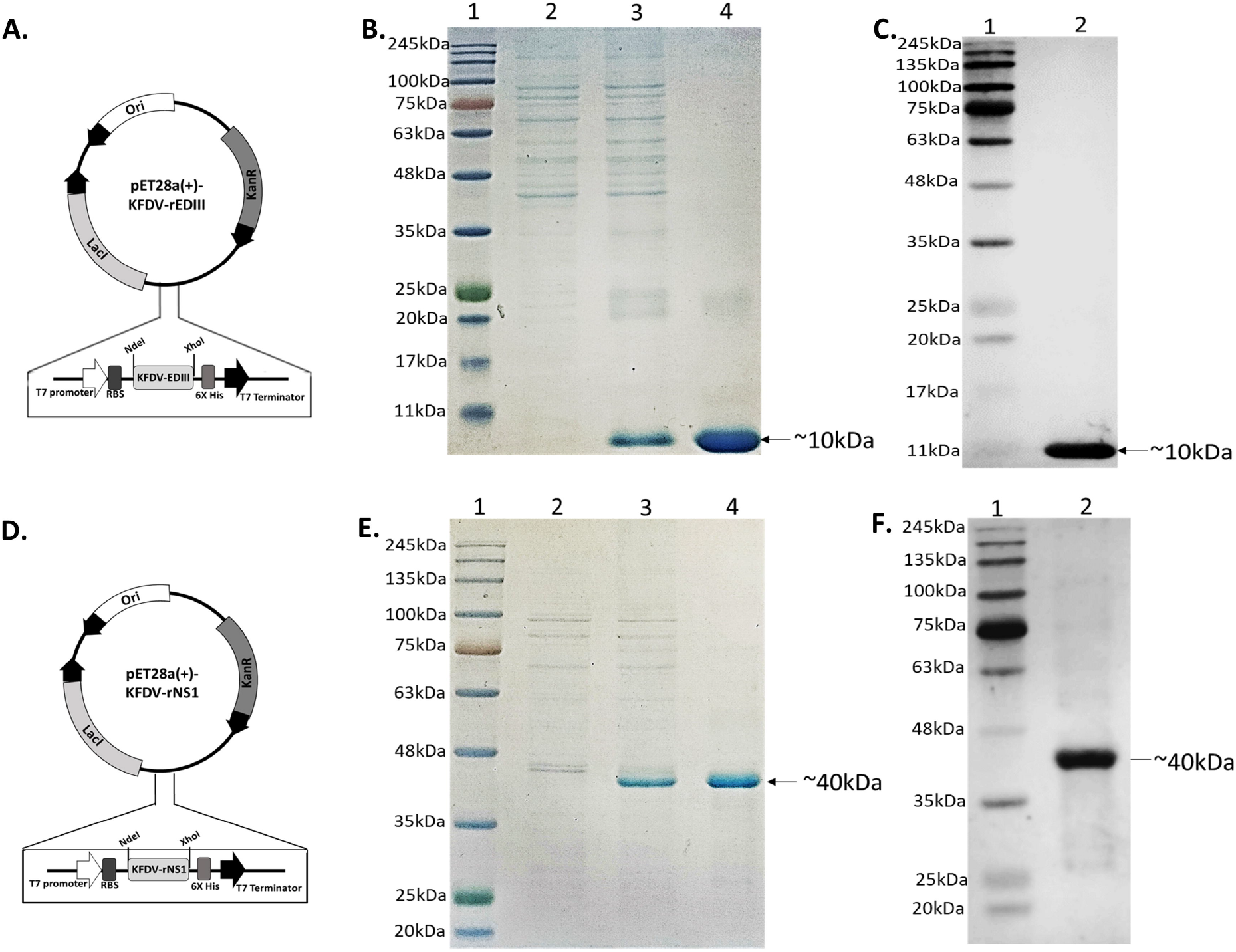
Expression profile of KFDV EDIII and NS1 proteins: The codon optimized sequences of KFDV EDIII and NS1 were inserted into bacterial expression pET28a(+) vector system. The pET28a(+)-KFDV-EDIII **(A)** and pET28a(+)-KFDV-NS1 **(D). B**, 15% SDS-PAGE analysis of KFDV-rEDIII protein expression, Lane 1: Pre-stained protein ladder (Himedia), Lane 2: *E. coli* lysate before IPTG induction, Lane 3: *E. coli* lysate after IPTG induction, Lane 4: Ni-NTA Affinity purified KFDV-rEDIII protein. **E**, 12 % SDS-PAGE analysis of rNS1 protein expression, Lane 1: Pre-stained protein ladder (Himedia), Lane 2: *E. coli* lysate before IPTG induction, Lane 3: *E. coli* lysate after IPTG induction, Lane 4: Ni-NTA Affinity purified rNS1 protein. Western blot analysis of KFDV marker proteins, rEDIII (**C)** (Lane 1: Protein ladder, Lane 2: purified KFDV-rEDIII probed using anti-his tag antibody), and rNS1 protein **(F)** (Lane 1: Protein ladder, Lane 2: purified KFDV-rNS1 probed using anti-his tag antibody).

### 3.2 Antibody titre evaluation of anti-KFDV-rEDIII and rNS1 polyclonal antibodies

The affinity-purified KFDV-rEDIII and rNS1 proteins were used to immunize female BALB/c mice and female New Zealand rabbits for the production of polyclonal antibodies. The antibody titres of anti-rEDIII and anti-rNS1 polyclonal antibodies were measured by indirect ELISA and dot-immunoblot assays to confirm immunogenicity. Initially, on the 14^th^ day, very low titres were observed for both the proteins; the end point titre (EPT) value paused at 1:16,000 for anti-rEDIII polyclonal antibodies and 1:8,000 for anti-rNS1 polyclonal antibodies. However, the titres increased significantly by 28^th^ and 42^nd^ day, particularly after the booster doses. By 70^th^ day, the EPT values were 1:256,000 for rEDIII and 1:512,000 for rNS1 (Figs. 2B and 2C). Additionally, the antibody titres for anti-rEDIII and rNS1 polyclonal antibodies were assessed using dot-blot assay. The titre values for these polyclonal antibodies were found to be 1:256,000 for rEDIII and 1:512,000 for the rNS1 protein (Figs. 2D and 2E). These findings indicate that both the recombinant KFDV proteins demonstrated high immunogenicity.

**Figure 2:**
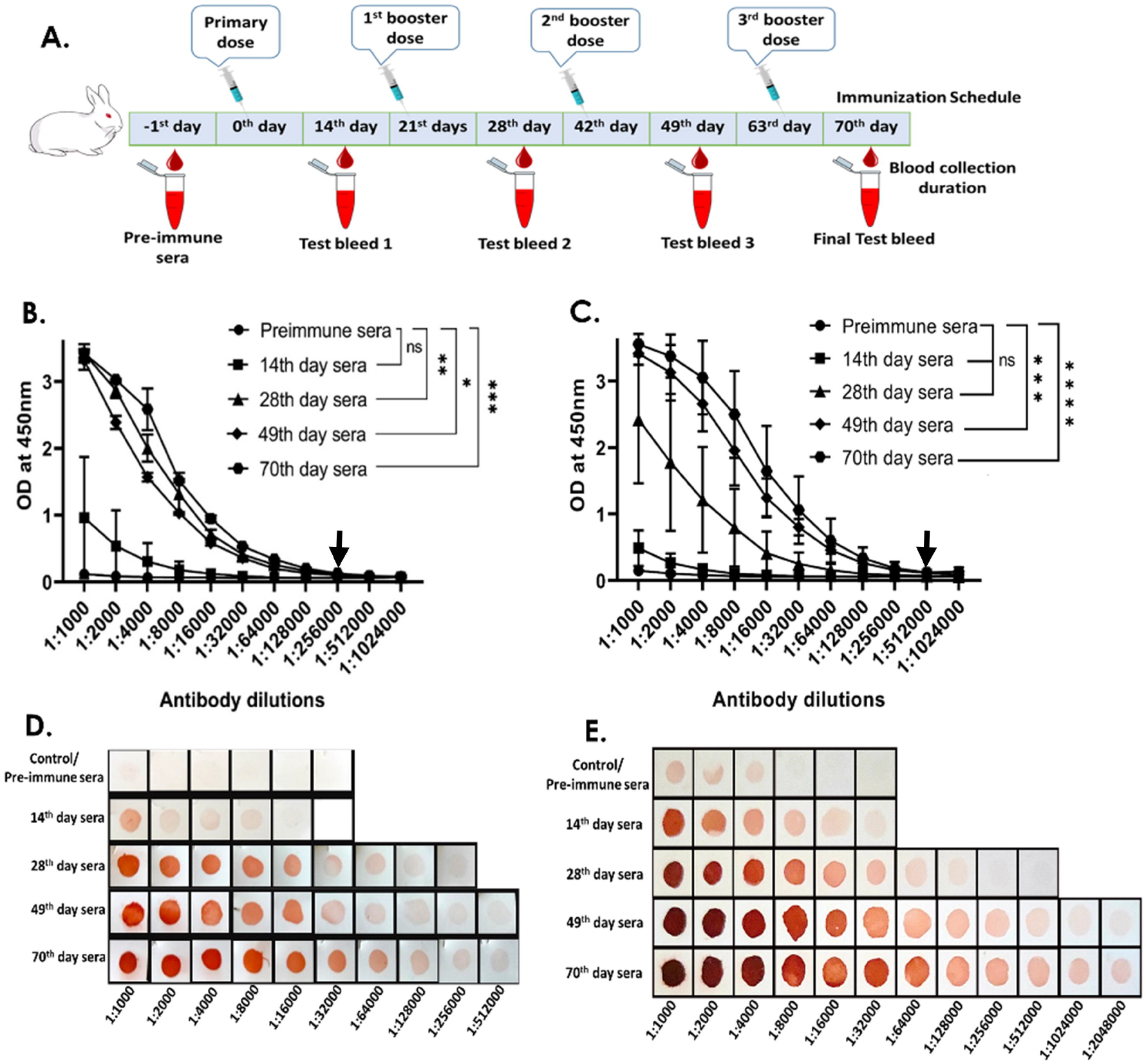
Immunogenicity and antibody titre evaluation of KFDV EDIII and NS1 proteins. **A**, Schematic representation of immunization and bleeding schedule. Immunogenicity analysis of KFDV-rEDIII protein (**B**) and rNS1 protein (**C**) by indirect ELISA. The data was statistically analysed using Graphpad Prism software version 8.3. The nested one-way ANOVA was utilised for the analysis and the results were expressed as * (*p*-value < 0.05), ** (*p*-value < 0.01), *** (*p*-value < 0.001) and ns (*nonsignificant*). **D & E**, Dot-blot Ab titre evaluation of anti-rEDIII and rNS1 polyclonal antibodies, respectively. The endpoint titre value (EPT) of anti-rEDIII polyclonal antibodies was 1:16,000 on 14^th^ day and 1:256,000 on 28^th^, 49^th^ and 70^th^ day. The EPT of anti-rNS1 polyclonal antibodies was 1:8,000 on 14^th^ day, 1:128,000 on 28^th^ day and 1:512,000 on 49^th^ and 70^th^ day). The final EPT value for anti-rEDIII and rNS1are indicated by an arrow.

### 3.3 Evaluation of cross-reactivity of anti-KFDV-rEDIII and rNS1 polyclonal antibodies against DENV positive samples and recombinant antigens

The KFDV-rEDIII and rNS1 proteins, along with their corresponding polyclonal antibodies, were evaluated for cross-reactivity with DENV using multiple criteria. Initially, a preliminary analysis was conducted using commercial dengue rapid detection kit. Dengue serum samples that tested positive for NS1 and IgM were used as positive controls, while PBS was used as a negative control. Both the KFDV recombinant proteins and the corresponding polyclonal antibodies showed no cross-reactivity (Figs. 3A and 3D). Subsequently, cross-reactivity was assessed using serum samples from individuals infected with DENV. A total of 41 DENV-positive serum samples were evaluated using the dengue rapid detection kit and categorized as either antigen positive or antibody positive (Supplementary Fig. S2). Out of 41 samples, 13 were positive for DENV-NS1 antigen, 16 were positive for DENV IgM/IgG antibodies, 4 samples were positive for both DENV-NS1 antigen and IgM/IgG antibodies, and 8 samples were tested negative for both DENV-NS1 antigen and IgM/IgG antibodies (Supplementary Table 1).

**Figure 3:**
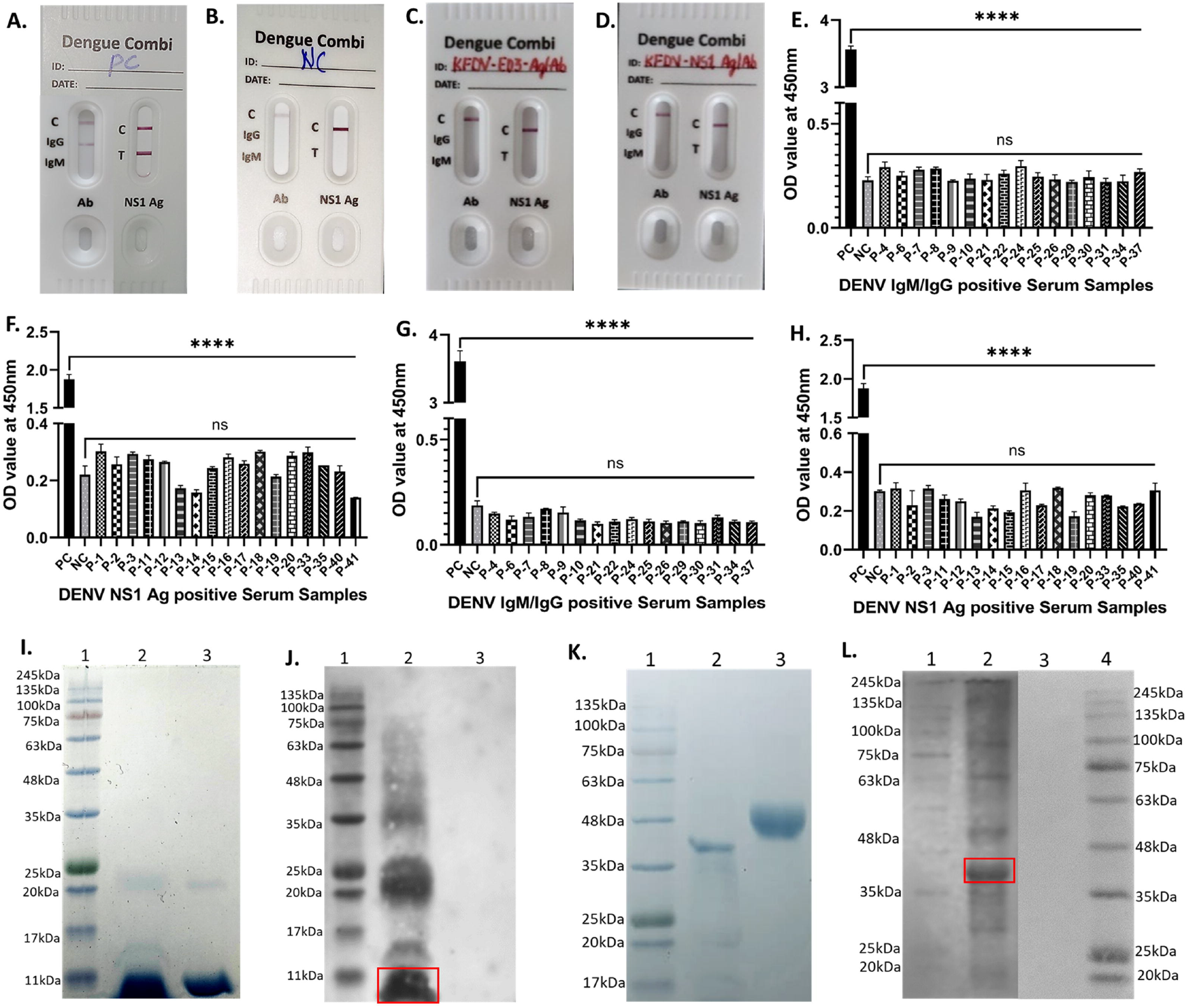
Cross-reactivity analysis of KFDV-rEDIII and rNS1 protein and antibody against DENV. Preliminary cross-reactivity analysis of bacterially expressed rKFDV antigens and anti-rKFDV polyclonal antibodies by using rapid DENV test kit (Agappe, India). **A**, Positive control (DENV antigen/antibody). **B**, Negative control (no antigen or unrelated protein). **C**, KFDV-rEDIII protein and anti-rEDIII antibody. **D**, KFDV-rNS1 protein and anti-rNS1 antibody. Antibody specificity analysis of dengue IgG/IgM positive samples against KFDV-rEDIII (**E**) and rNS1 protein (**G**). Antibody cross-reactivity analysis of anti KFDV-rEDIII (**F**) and rNS1 against DENV-NS1 antigen positive samples (**H**). The data were analysed using Graph Pad prism (v8.3). Nested one-way ANOVA was used for the analysis of data and the results were expressed as ns, non-significant, and **** (p<0.0001). **I**, 15% SDS-PAGE profile of KFDV–rEDIII and DENV-rEDIII protein. Lane 1: Protein ladder, Lane 2: purified KFDV-rEDIII protein, Lane 3: DENV-rEDIII protein (Native Antigen, UK). **J**, Western blot analysis of cross-reactivity of anti KFDV-rEDIII antibodies against DENV-rEDIII protein. Lane 1: Protein ladder, Lane 2: KFDV-rEDIII protein, Lane 3: DENV-rEDIII protein (Native Antigen, UK). **K**, 12% SDS-PAGE profile of KFDV-rNS1 and DENV-rNS1 protein. Lane 1: Protein ladder, Lane 2: purified KFDV-rNS1 protein, Lane 3: DENV-rNS1 protein (Native Antigen, UK). **L**, Western blot analysis of cross-reactivity of anti KFDV-rNS1 antibodies against DENV-rNS1 protein. Lane 1: Protein ladder, Lane 2: KFDV-rNS1 protein, Lane 3: DENV-rNS1 protein (Native Antigen, UK), Lane 4: Protein ladder.

The dengue IgM/IgG positive samples were analysed for antibody specificity against recombinant KFDV proteins through indirect ELISA. In this assay, both rKFDV proteins were used as antigens for DENV IgM/IgG positive sera. Anti-KFDV-rEDIII and rNS1 polyclonal antibodies were employed as positive controls, while serum from healthy uninfected individuals was used as a negative control. The DENV IgM/IgG positive serum did not detect either of the KFDV recombinant proteins (rEDIII and rNS1), indicating no cross-reactivity between DENV IgM/IgG antibodies and KFDV recombinant proteins (Figs. 3B and 3E). For antigen specificity analysis, DENV-NS1 antigen positive samples were tested against anti-rEDIII and rNS1 polyclonal antibodies by using DENV-NS1 sandwich ELISA commercial kit. This kit included anti-DENV-NS1 antibody pre-coated onto micro-wells, along with positive and negative control samples. The DENV-NS1 antigen positive samples were incubated with anti-rEDIII and rNS1 polyclonal antibodies. None of the anti-rEDIII or rNS1 polyclonal antibodies detected DENV-NS1 antigen, indicating the absence of cross-reactivity (Figs. 3C and 3F).

In addition, the cross-reactivity between KFDV polyclonal antibodies and DENV antigens was verified by western blotting. Recombinant DENV antigens (rEDIII and rNS1) were analyzed using anti-KFDV-rEDIII and rNS1 polyclonal antibodies, with KFDV-rEDIII and rNS1 proteins serving as positive controls (Figs. 3G and 3I). The assay confirmed no cross-reactivity, and only the recombinant KFDV proteins were detected, and not recombinant DENV proteins (Figs. 3H and 3J).

### 3.4 Detection of specificity and sensitivity of KFDV specific antigen-based sandwich ELISA

We have successfully developed a cutting-edge antigen-based sandwich ELISA for the accurate KFDV antigen detection. Utilization of mice polyclonal capture antibodies (1:50), coupled with rabbit-derived polyclonal antibodies for detection (1:1000), this novel assay offers a highly specific method for KFD antigen identification. The sandwich ELISA detected KFDV-rEDIII antigen at a very low concentration of 3ng and the KFDV-rNS1 antigen at a concentration of 25ng in PBS (Figs. 4A and 4D). The detection ability of this sandwich ELISA decreased when these recombinant proteins were spiked into human serum to the level of 300ng for KFDV-rEDIII and 2µg for KFDV-rNS1 (Figs. 4B and 4E). Further, it is clear that the anti-KFDV-rEDIII antibodies exhibited a heightened sensitivity as compared to their anti-KFDV-rNS1 counterparts. Moreover, the sandwich ELISA for KFDV-rEDIII and rNS1 polyclonal capture and detection antibodies were highly specific, exhibiting no cross-reactivity with DENV-rEDIII and rNS1 proteins (Figs. 4C and 4F).

**Figure 4:**
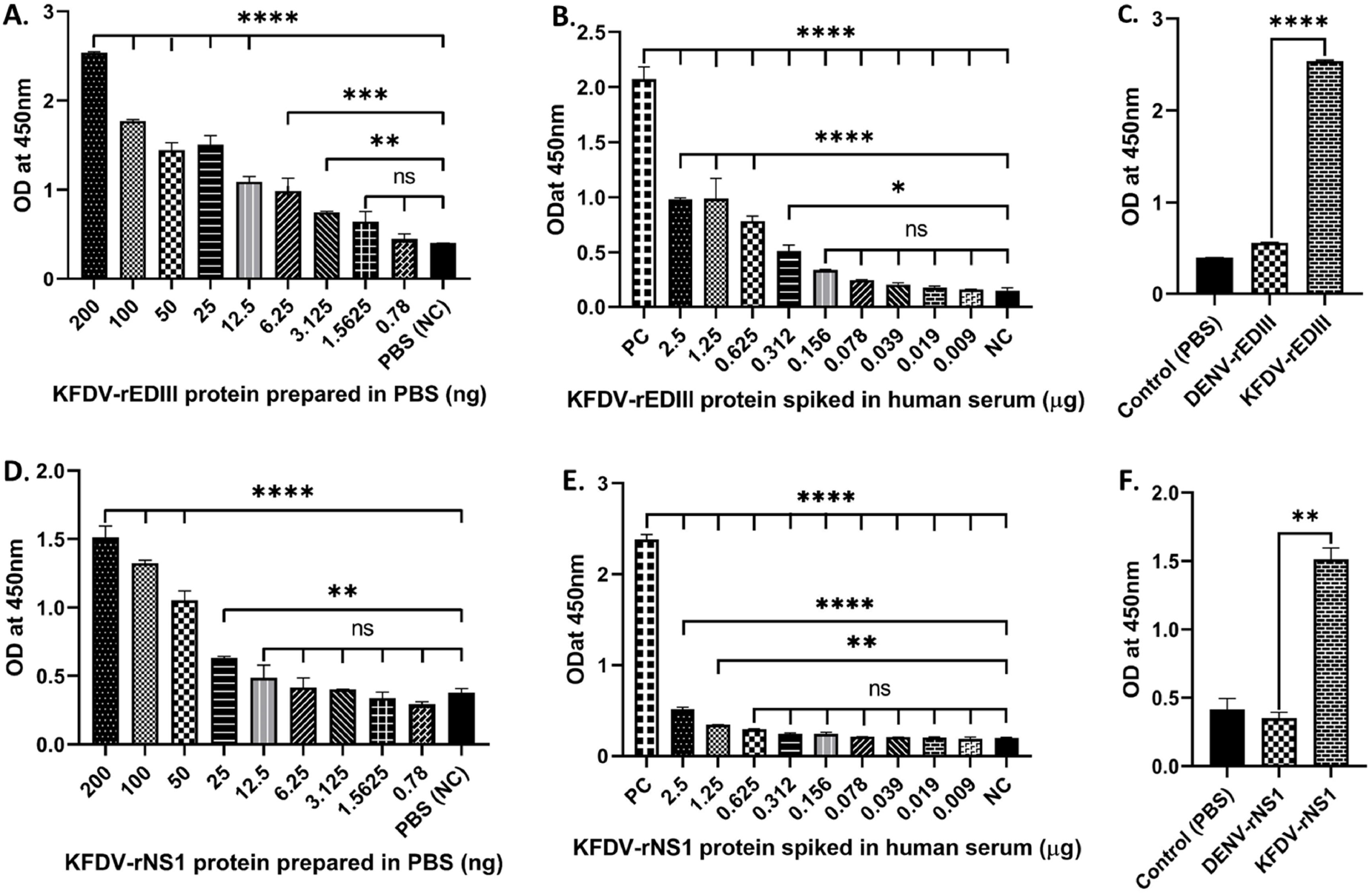
Sensitivity and specificity analysis of KFDV specific sandwich ELISA: Sensitivity analysis of this sandwich ELISA was tested against different dilutions of KFDV-rEDIII protein (**A**) and rNS1 protein (**D**) in PBS and after spiking in human serum, KFDV-rEDIII protein (**B**) and rNS1 protein (**E**). The specificity analysis of DENV-rEDIII and KFDV-rEDIII protein (**C**) and DENV-rNS1 and KFDV-rNS1 protein (**F**). Data was analysed using one-way ANOVA in Graphpad Prism version 8.3. The results were expressed as * (p < 0.05), ** (p < 0.01), *** (p < 0.001), **** (p < 0.0001) and *ns* (nonsignificant).

### 3.5 Polyclonal antibodies raised against bacterially expressed KFDV-rEDIII specifically recognize KFDV E protein expressed in mammalian cells

In the sensitivity analysis of KFD specific sandwich ELISA, we found that only the KFDV-rEDIII protein was detected after spiking in the serum. Therefore, we evaluated the detection capability of anti-rEDIII and rNS1 polyclonal antibodies against full-length glycosylated KFDV E protein. The HEK293 cells were first transfected with the pCDNA3.1(+)-KFDV-E vector, resulting in stable HEK293 cells expressing full-length KFDV E protein (Fig. 5A). The polyclonal antibodies raised against bacterially-expressed KFDV-rEDIII successfully detected the mammalian-expressed complete KFDV E protein (Fig. 5B, i). In contrast, the polyclonal antibodies raised against the bacterially-expressed KFDV-rNS1 did not detect the mammalian-expressed complete KFDV E protein (Fig. 5B, ii) pointing no cross-reactivity between anti-rEDIII and rNS1 polyclonal antibodies.

**Figure 5:**
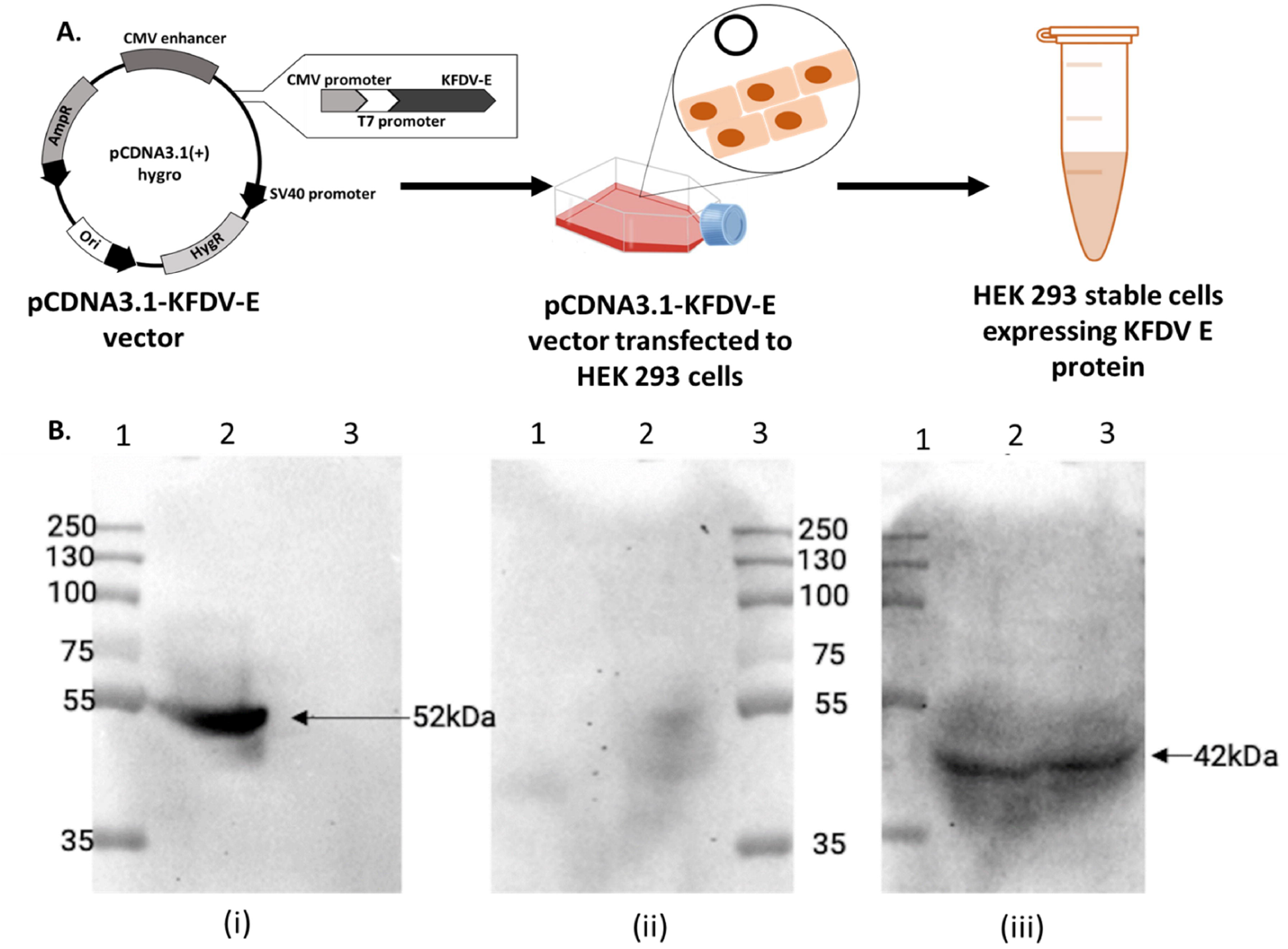
Validation of polyclonal antibodies against glycosylated KFDV E protein. **A**, Schematic representation of KFDV E protein (complete) construct in pCDNA3.1 (+) Hygro vector and stable protein expression in HEK293 mammalian cells. **B**, Evaluation of polyclonal antibody specificity against mammalian expressed KFDV E protein by Western blot analysis. **B.i**, HEK293-KFDV E protein against anti-KFDV-rEDIII polyclonal antibody (Lane 1, protein ladder; Lane 2, stable lysate of HEK293 expressing KFDV E protein; Lane 3, mock HEK293 lysate). **B.ii**, HEK293-KFDV E protein against anti-KFDV-rNS1 polyclonal antibody (Lane 1, mock HEK293 lysate; Lane 2, stable lysate of HEK293 expressing KFDV E protein; Lane 3, protein ladder). **B.iii**, HEK293-KFDV E protein against reference housekeeping protein β-actin antibodies (Lane 1, protein ladder; Lane 2, stable lysate of HEK293 expressing KFDV E protein; Lane 3, mock HEK293 lysate).

## 4. Discussion

KFD is a neglected tropical disease mostly prevalent in Southern India. The disease was once endemic only to Karnataka state but is now showing its presence in other regions of Western Ghat, extending from Kerala to Gujarat (21,22). Currently, there are no effective treatment strategies for KFD. Although a poorly immunogenic formalin-inactivated KFD vaccine was used until 2022 in the endemic areas (23), its widespread refusal is primarily due to the irritation caused by residual formalin contaminants in the vaccine (24). Consequently, there is an increasing emphasis on effective newer KFD vaccines, by exploring highly immunogenic viral vector-based platforms (25,26). Moreover, accurate rapid diagnostic methods are crucial for managing infectious diseases, especially in the context of disease outbreaks in remote areas. Current KFD diagnosis involves combination of two techniques, in which KFD-specific IgM ELISA has shown cross-reactivity with other viruses such as DENV and chikungunya (CHIKV), whereas RT-PCR is highly specific but requires sophisticated facilities (9,27).

In view of the above, our objective was to develop an antigen-based diagnostic to serve as an efficient method for quick and accurate field detection of KFD. This is particularly important in endemic areas to differentiate KFD from other cross-reactive arboviral diseases (28). To achieve this, we targeted the KFDV-EDIII and KFDV-NS1 proteins as biomarkers. These flaviviral proteins have been proven to be prominent serum biomarkers and potential vaccine candidates (10,29). Initially, we used bacterial expression system to obtain these proteins in high yields. However, the expressed proteins were found to be insoluble because of their presence in the inclusion bodies, necessitating a renaturation process after purification (30). The recombinant NS1 protein expressed in bacteria has been extensively utilized in diagnosing DENV, demonstrating its effectiveness over NS1 proteins expressed in mammalian systems (31,32). The data also suggests that protein glycosylation does not interfere with dengue diagnosis which means, the proteins expressed in bacterial system are also acceptable for immunological studies although they may lack conformational antigenic determinants. Further, refolded bacterial-expressed proteins can be effectively used for vaccine development and diagnostic purposes, as they retain the native structural conformation (33,34). Additionally, mammalian and insect cell-based expression methods are laborious and yield lower amounts of the desired protein. Thus, bacterial expression and proper refolding of protein, addresses these issues indicating its utility.

The immunogenicity of expressed proteins was further evaluated by generating polyclonal antibodies in rabbits and assessed their reactivity by indirect ELISA and immunoblotting. These assays demonstrated that both rEDIII and rNS1 proteins are immunogenic and capable of eliciting a strong immune response, as indicated by high antibody titres, which is crucial for an effective and sensitive diagnostic tool. The flaviviral EDIII and NS1 proteins were even considered for the vaccine development resulting in high neutralising antibody titre as well as effective T-cell activation (35,36).

A highly conserved protein epitope that induces cross-protection could be valuable for the development of broad range protective vaccines as the antibodies generated can cross-neutralize the pathogen and can also activate memory CD4 cells (37,38). In contrast, when developing specific diagnostic assays, the highly distinct epitopes with low or less conserved sequences should be considered as they yield only non-cross reactive antibodies (39). Notably, in our study, the polyclonal antibodies generated against EDIII and NS1 proteins were highly specific, with no cross-reactivity with other flaviviruses such as DENV. This specificity is a key advantage for KFD diagnostics, as flaviviral cross-reactivity often presents a major challenge in the accurate detection of KFD, particularly in multiple flavivirus infection endemic regions (27). The observed non-cross reactivity suggests that these antibodies can serve as reliable markers for distinguishing KFDV infections from other tick-borne or mosquito-borne viruses.

This study also involved the development of a sandwich ELISA using polyclonal antibodies generated against KFD-EDIII and NS1 proteins. The resulting KFD-specific sandwich ELISA, developed using anti-rEDIII polyclonal antibodies demonstrated high sensitivity and specificity, aiding the detection of KFDV antigens in serum samples and this approach has advantages. Firstly, the sandwich format enhances the sensitivity by amplifying the interaction through the use of two different antibodies targeting distinct epitopes on the same antigen. The sensitivity of this method matches that of the dengue NS1-specific ELISA, as the concentration of NS1 among infected individuals ranges from 10 ng to 2 µg during active infection (40). Secondly, targeting KFDV specific antigens rather than the antibodies, minimizes the potential for false positives, which is a common concern with other flaviviral diagnostic tests that might cross-react with DENV or CHIKV antibodies. This method was able to specifically detect KFDV-EDIII antigen which makes this as a precise assay as well as potential tool for early KFD detection, ultimately contributing to better disease management and control.

A recent study was aimed at improving the diagnosis of KFD by IgM-ELISA using KFDV EDIII antibodies that are produced after the infection (41). However, their focus was to detect IgM antibodies generated against KFDV-EDIII rather than EDIII or E antigen itself. Further, the problem of cross-reactivity may potentially persist here, as seen in the case of DENV. For instance, DENV detection employ EDIII-specific IgM antibodies to detect secondary DENV infections, which can lead to potential cross-reactivity (42,43). In contrast, our study focussed on detecting the antigen which targets only active infections as the current KFD diagnostic methods target antibodies which could potentially identify the past infections also and thus may not be suitable for early-stage diagnosis. By detecting KFDV antigens in the serum, the sandwich ELISA offers a tool for the early KFD diagnosis. This can help to improve treatment regime and enable more effective disease outbreak management.

Another study conducted by Yadav et al. (44) utilized the polyclonal antibodies of KFDV-NS1 protein for the early detection of KFD. This study has not provided the supportive experimental evidence regarding antibody titers, cross-reactivity, or ROC (Receiver Operating Characteristic) data, apart from the expression pattern of KFDV-NS1 protein. However, they reported that these antibodies failed to detect KFDV-NS1 antigens in the serum samples. This could be attributed to the degradation of NS1 protein in KFD patient serum, as the timing of sample collection plays a crucial role in early detection. Typically, the flaviviral NS1 protein is released into the bloodstream soon after infection, approximately within hours. Therefore, many DENV detection kits incorporate both NS1 antigen as an early marker and anti-IgM/IgG antibodies to identify secondary infections (45,46,29). In contrast to the studies conducted by Yadav et al. (44), our study addressed all relevant aspects with experimental support, except the validation using KFD-positive samples through ROC curve plotting. Although we did not validate the assay using actual KFD infected samples, we conducted a serum spike assay to mimic natural infection, in line with a previous study that utilized antibody-spiked serum (47). The results indicate that the KFDV-rEDIII specific polyclonal antibodies demonstrate higher sensitivity, as it can detect the rEDIII antigen even at low levels (300 ng) in the serum. Envelope protein is highly stable and plays a crucial role in maintaining the structural integrity of the virus, which does not have any role in viral replication and have fewer cleavage sites (48). Because of these characteristics, the envelope protein is not significantly affected by serum proteases. However, rNS1 anti-rabbit polyclonal antibodies have nearly failed to detect the rNS1 protein when spiked into human serum, which needs to be validated with actual KFD-infected serum samples. This issue may stem from the intrinsic properties of the NS1 protein rather than its antibodies. NS1 is known to be highly unstable and exists in multiple oligomeric forms. Notably, it circulates in serum predominantly as a soluble hexamer (sNS1), which plays a role in immune evasion, while the dimeric form localized in the cytoplasm is associated with viral replication (49). We further speculate that NS1 may possess multiple enzymatic or proteolytic cleavage sites, making it susceptible to degradation by serum proteases. In this context, the hexameric configuration could confer structural protection against proteolysis (50). The decreased sensitivity observed with antigen-spiked serum can be attributed to the matrix effect, where endogenous serum components such as proteins, lipids, and surfactants interfere with the antigen–antibody binding or increase background noise, thereby reducing the overall detection efficiency of the assay.

Polyclonal antibodies raised against bacterial EDIII successfully detected the full-length glycosylated E protein of KFDV and did not show cross-reactivity with rabbit anti-rNS1 polyclonal antibodies. These results demonstrate that the bacterial-expressed protein can be used as an unambiguous diagnostic marker. Overall, we have successfully developed a sandwich ELISA that is very specific and sensitive. However, the sensitivity may not be higher than the currently used RT-PCR, but the proposed method benefits the early detection.

## Limitations of the study

In this study, although we successfully developed a specific KFD diagnostic assay, there are some limitations. One major limitation is that the assay needs to be validated with a larger and more diverse sample sets (active virus culture), including clinical samples from KFD infected patients. Additionally, it is crucial to investigate cross-reactivity with other closely related flaviviruses which will ensure the test robustness in endemic regions. Another aspect that requires improvement is the assay’s sensitivity in detecting low viral loads, which is vital for diagnosing early-stage or asymptomatic cases of KFD. To enhance the performance of the assay, one could consider using monoclonal antibodies or developing quantitative sandwich ELISA.

## 5. Conclusions

In this study, we successfully developed a KFD-specific sandwich ELISA using bacterially expressed KFDV-EDIII, offering a promising tool for rapid, specific and sensitive diagnosis of KFD. The non-cross reactive polyclonal antibodies used in this study represents a significant advancement in KFD diagnosis, effectively addressing many challenges associated with cross-reactivity in flavivirus diagnostics. This diagnostic assay lays the foundation for future research aimed at optimizing the test for clinical applications and assessing its performance in field conditions. Finally, integrating this diagnostic tool into the disease surveillance programs with point-of-care diagnosis could enhance the early detection, monitoring, management, and control of KFD outbreaks, especially in endemic regions. Further the use of anti-KFDV-rEDIII monoclonal antibodies could enhance the sensitivity and specificity of the existing method suitable for on-field detection.

## Data availability statement

All data is available within the manuscript and supplementary files.

## Author contribution statement

Sayad Hafeez involved in conducting the experiments and manuscript writing. N.B. Thippeswamy conceptualized the study, supervised the work, and corrected the manuscript. Rajeshwara N. Achur helped in the study design and data interpretation. Easwaran Sreekumar performed the mammalian expression based experiments. Suchetha Kumari and Asha Srinivasan involved in sample collection and cross reactivity assays.

## Acknowledgements

The authors are thankful to VGST, Karnataka, India, (GRD No.1119) for the financial assistance provided in conducting this study and Lady Tata Memorial Trust, Mumbai, India, for awarding Senior Research Fellowship to Mr. Sayad Hafeez (2020-2023). Authors extend the gratitude towards Chromed biosciences Pvt. Ltd., Tumakuru, for allowing to use their animal facility. The authors acknowledge Mr. Vivek Vijay, Project Associate, Molecular Bioassay Laboratory, Institute of Advanced Virology (IAV), Thiruvananthapuram, Kerala, for experimental support with the mammalian cell expression based western blots.

