## Supplemental Fig. S1, S2 and Table 1 for "Utility of recombinant envelope domain III as a diagnostic antigen for the specific detection of Kyasanur Forest Disease"

**Unequivocal detection of Kyasanur Forest Disease virus using polyclonal antibodies against recombinant EDIII and NS1 proteins**

**Supplementary Information:**

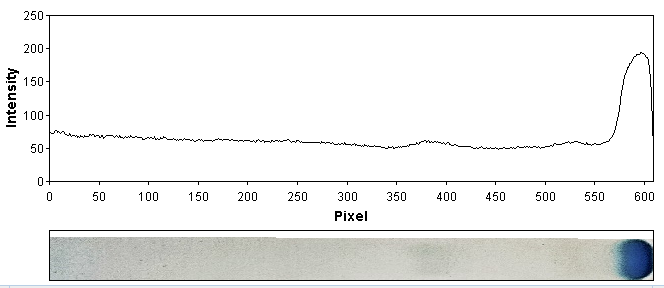

**A.**

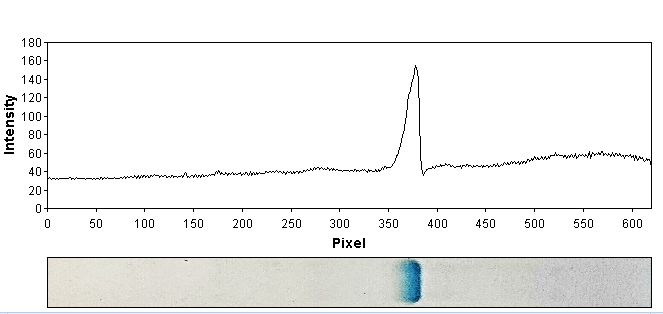

**B.**

**Figure S1: Densitometry analysis of purified rKFDV proteins**. **A**, rEDIII protein and **B**, rNS1 protein.

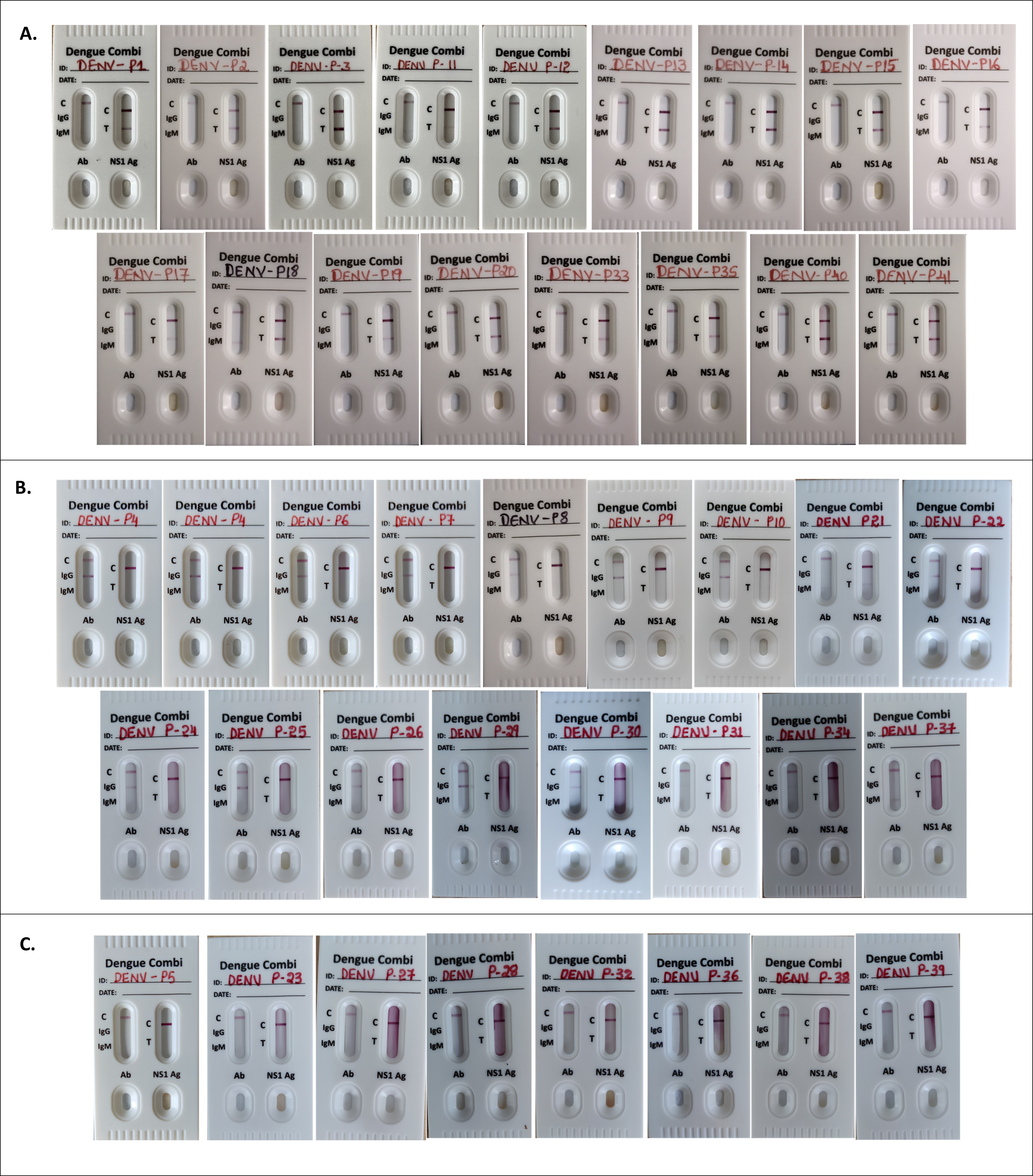

**Figure S2: Cross-verification of collected DENV serum samples.** DENV positive serum samples collected from district hospitals were cross-verified using rapid DENV detection kit (Agappe dianostics ltd., Kerala, India). Out of 41 serum samples 17 samples were positive for DENV NS1 (A), 17 samples were positive for IgM/IgG (B) and 8 samples were negative (C).

**Supplementary Table 1:** Details of DENV positive samples collected from the K.S Hegde Medical hospital, Mangaluru.

| **Sl. No.** | **Patient ID** | **Dengue NS1 ELISA** | **Dengue IgM/IgG ELISA** | **Dengue Kit analysis** | |
| --- | --- | --- | --- | --- | --- |
|  |  |  |  | **NS1 positive** | **IgM/IgG positive** |
| 1. | P1 | **+** | **-** | **+** | **-** |
| 2. | P2 | **+** | **-** | **+** | **-** |
| 3. | P3 | **+** | **-** | **+** | **-** |
| 4. | P4 | **-** | **+** | **-** | **+** |
| 5. | P5 | **-** | **-** | **-** | **-** |
| 6. | P6 | **-** | **+** | **-** | **+** |
| 7. | P7 | **-** | **+** | **-** | **+** |
| 8. | P8 | **-** | **+** | **-** | **+** |
| 9. | P9 | **-** | **+** | **-** | **+** |
| 10. | P10 | **-** | **+** | **-** | **+** |
| 11. | P11 | **+** | **-** | **+** | **+** |
| 12. | P12 | **+** | **-** | **+** | **-** |
| 13. | P13 | **+** | **-** | **+** | **-** |
| 14. | P14 | **+** | **-** | **+** | **-** |
| 15. | P15 | **+** | **-** | **+** | **-** |
| 16. | P16 | **+** | **-** | **+** | **-** |
| 17. | P17 | **+** | **-** | **+** | **-** |
| 18. | P18 | **+** | **-** | **+** | **+** |
| 19. | P19 | **+** | **-** | **+** | **-** |
| 20. | P20 | **+** | **-** | **+** | **-** |
| 21. | P21 | **-** | **+** | **-** | **+** |
| 22. | P22 | **-** | **+** | **-** | **+** |
| 23. | P23 | **-** | **-** | **-** | **-** |
| 24. | P24 | **-** | **+** | **-** | **+** |
| 25. | P25 | **-** | **+** | **-** | **+** |
| 26. | P26 | **-** | **+** | **-** | **+** |
| 27. | P27 | **-** | **-** | **-** | **-** |
| 28. | P28 | **-** | **-** | **-** | **-** |
| 29. | P29 | **-** | **+** | **-** | **+** |
| 30. | P30 | **-** | **+** | **-** | **+** |
| 31. | P31 | **-** | **+** | **-** | **+** |
| 32. | P32 | **-** | **-** | **-** | **-** |
| 33. | P33 | **+** | **-** | **+** | **-** |
| 34. | P34 | **-** | **+** | **-** | **+** |
| 35. | P35 | **+** | **-** | **+** | **+** |
| 36. | P36 | **-** | **-** | **-** | **-** |
| 37. | P37 | **-** | **+** | **-** | **+** |
| 38. | P38 | **-** | **-** | **-** | **-** |
| 39. | P39 | **-** | **-** | **-** | **-** |
| 40. | P40 | **+** | **-** | **+** | **-** |
| 41. | P41 | **+** | **-** | **+** | **+** |
